# Neural speech tracking shifts from the syllabic to the modulation rate of speech as intelligibility decreases

**DOI:** 10.1101/2021.03.25.437033

**Authors:** Fabian Schmidt, Ya-Ping Chen, Anne Keitel, Sebastian Rösch, Ronny Hannemann, Maja Serman, Anne Hauswald, Nathan Weisz

**Affiliations:** Centre for Cognitive Neuroscience, University of Salzburg, 5020 Salzburg, Austria; Department of Psychology, University of Salzburg, 5020 Salzburg, Austria; Psychology, School of Social Sciences, University of Dundee, DD1 4HN Dundee, UK; Department of Otorhinolaryngology, Paracelsus Medical University, 5020 Salzburg, Austria; Audiological Research Unit, Sivantos GmbH, 91058 Erlangen, Germany

## Abstract

The most prominent acoustic features in speech are intensity modulations, represented by the amplitude envelope of speech. Synchronization of neural activity with these modulations is vital for speech comprehension. As the acoustic modulation of speech is related to the production of syllables, investigations of neural speech tracking rarely distinguish between lower-level acoustic (envelope modulation) and higher-level linguistic (syllable rate) information. Here we manipulated speech intelligibility using noise-vocoded speech and investigated the spectral dynamics of neural speech processing, across two studies at cortical and subcortical levels of the auditory hierarchy, using magnetoencephalography. Overall, cortical regions mostly track the syllable rate, whereas subcortical regions track the acoustic envelope. Furthermore, with less intelligible speech, tracking of the modulation rate becomes more dominant. Our study highlights the importance of distinguishing between envelope modulation and syllable rate and provides novel possibilities to better understand differences between auditory processing and speech/language processing disorders.

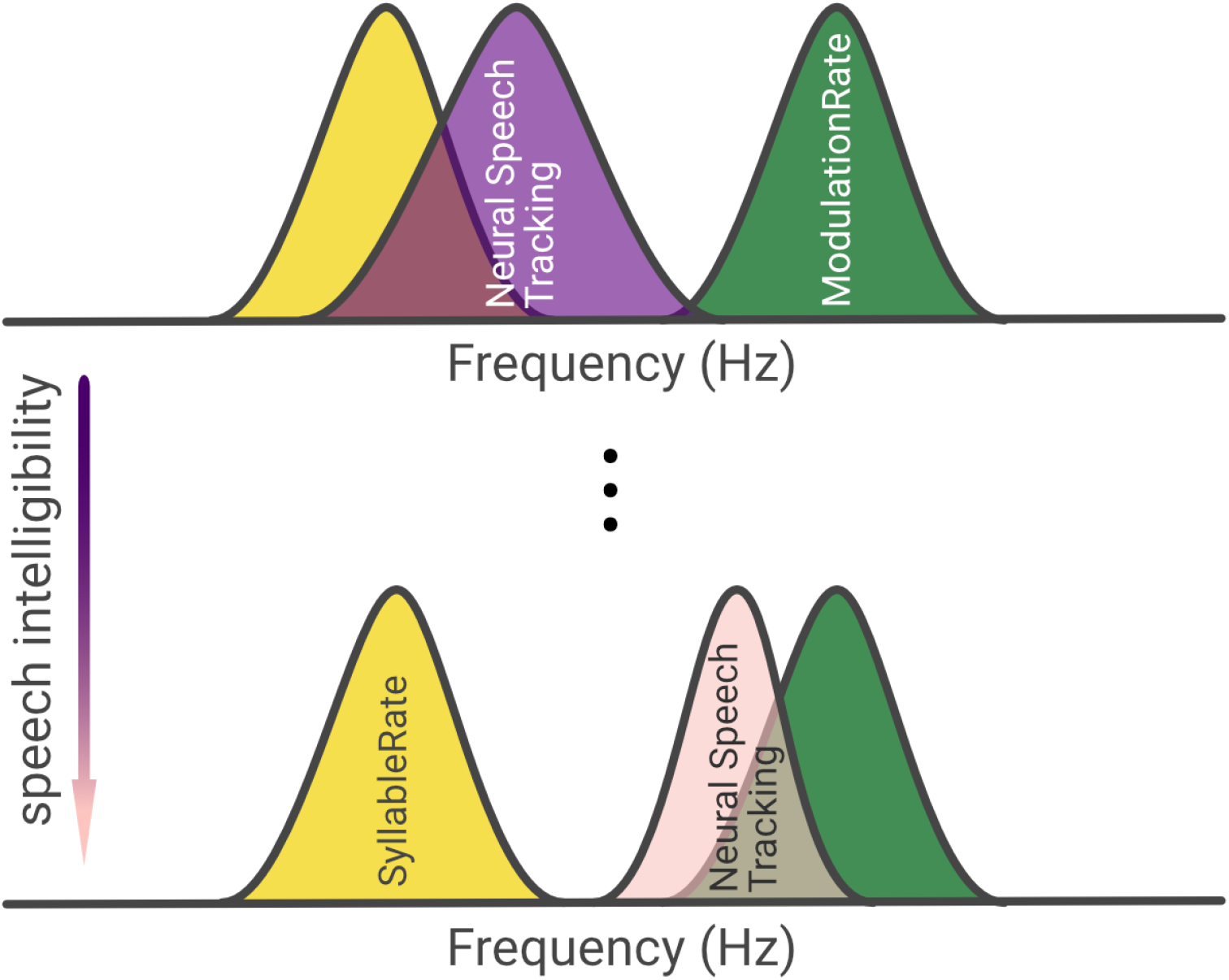

## INTRODUCTION

Intensity modulations of the acoustic envelope reflect the most prominent feature of the acoustic speech stream. Synchronization of neural activity with these modulations is vital for speech comprehension (*1*–*4*). As the acoustic modulation of speech and the production of syllables is correlated (*5*), investigations of neural speech tracking rarely distinguish between the lower-level acoustic and higher-level linguistic information. However, while the temporal scale of the acoustic modulation (∼4-5 Hz) is remarkably similar across languages, speakers, and speaking conditions (for reviews see *5, 6*), the rate at which syllables are produced can vary significantly across (and within) languages (*7*), dialects and speaking conditions (*8*). Therefore, it remains unclear whether and how the brain differentially tracks low-level acoustic and linguistic information during natural continuous speech. Distinguishing these aspects more clearly may also be important in gaining a better understanding of the neural processes separating auditory processing disorders (e.g. hearing loss) from language processing disorders (e.g. developmental dyslexia).

The ability to process meaningful information from an acoustic sound stream becomes especially important in difficult listening situations. While some studies indicate a positive relationship between speech intelligibility and the synchronization of brain activity with the speech envelope (neural speech tracking) in the low-frequency range (*1*–*4*) others have reported inverse effects (*9, 10*). A recent study even suggested an inverted u-shaped relationship, where synchronization increases when speech is mildly degraded and decreases as speech becomes unintelligible (*11*). This wide range of (partly contradicting) results is suggestive of a complex relationship between the intelligibility of speech and the related neural dynamics of speech tracking.

One source of these seeming inconsistencies may be caused by the interpretation of band-limited differences, conflating periodic (center frequency, power, bandwidth) and aperiodic (offset, exponent) properties of the underlying signal (*12*). In fact, both the acoustic envelope of speech and electrophysiological measurements of neural activity possess an overall 1/f-like spectrum (*13, 14*). This 1/f-like pattern is also at times present in the low-frequency coherence/correlation spectrum between both signals (e.g. see *1, 9, 11*). Recently, several approaches were proposed to separate periodic from aperiodic components of electrophysiological activity (IRASA (*15*); FOOOF (*12*)). We applied one of these approaches (FOOOF) to speech tracking, to parametrize the periodic components underlying low-frequency speech-brain coherence, such as the center frequency, the relative height of the coherence peak, and its bandwidth (∼tuning). Commonly, when investigating neural speech tracking these parameters are not separated from the aperiodic components of the coherence spectra. Instead (band/-averaged) contrasts over coherence spectra across several experimental conditions are computed, conflating the periodic and aperiodic components underlying speech-brain coherence. We propose that the periodic components (center frequency, relative height of the coherence peak, bandwidth) of speech-brain coherence offer a better estimate of neural speech tracking than broadband speech-brain coherence in the conventional frequency ranges. Therefore, it may be beneficial to investigate these parameters separately to better understand how neural activity tracks lower-level and higher-level information in a continuous speech stream and how this tracking is influenced by speech intelligibility.

Here, we applied this approach to two separate studies in which speech intelligibility was parametrically controlled via vocoding (3-, 7-Channels or no vocoding). Vocoding (*16*) is a popular technique to manipulate the intelligibility of speech that allows for high parametric control, while only moderately influencing the acoustic envelope of the signal (*2*). We captured the spectral dynamics of neural speech processing at cortical and subcortical levels (*17*) of the auditory hierarchy using magnetoencephalography (MEG). We observed that low-frequency speech brain coherence in accordance with previous results (*1*–*4*) declines with a decrease in intelligibility. However, parametrization of the coherence spectra revealed that this effect was mainly driven by the aperiodic components. The periodic components that are actually reflective of neural speech tracking (as opposed to band-limited coherence differences) were characterized by a narrower frequency tuning of the low-frequency coherence peak of vocoded speech along with an increase of its center frequency. The latter effect points to a shift of cortical tracking away from the syllabic rate towards the modulation rate of the acoustic envelope as vocoding increased. This effect is also seen for subcortical regions, although tracking is here overall dominated by the modulation rate.

## RESULTS

### Task performance declines with speech intelligibility

Subjects (*N*=55 across 2 experiments; Fig. 1B,C) listened to an audiobook (“Das Märchen”; Goethe, 1795) narrated by a female speaker whilst seated in the MEG. Parts of the audiobook presented were noise-vocoded (Fig. 1A; 7-Chan, 3-Chan). Vocoding levels were either kept constant throughout the audio presentation (Study#1; Fig. 1B) or changed intermittently (Study#2; Fig. 1C) to test the influences of vocoded speech on neural speech tracking under two different conditions. At the end of each audio presentation, subjects were presented with two nouns from which they had to pick the one they perceived in the previous sentence. The audio presentations were embedded in blocks that varied between 3.5 and 9 minutes (see Methods & Materials for a detailed account). Due to the overall low number of behavioral responses, we added an additional behavioral assessment (adjusted for each study) to investigate how vocoding influences speech comprehension. The task was similar to the one performed in the actual measurement but consisted of a larger amount of shorter trials (24; see Materials & Methods for a detailed account). Due to technical difficulties, only a subset (*N*=39) of our subjects participated in these assessments.

**Fig 1.**
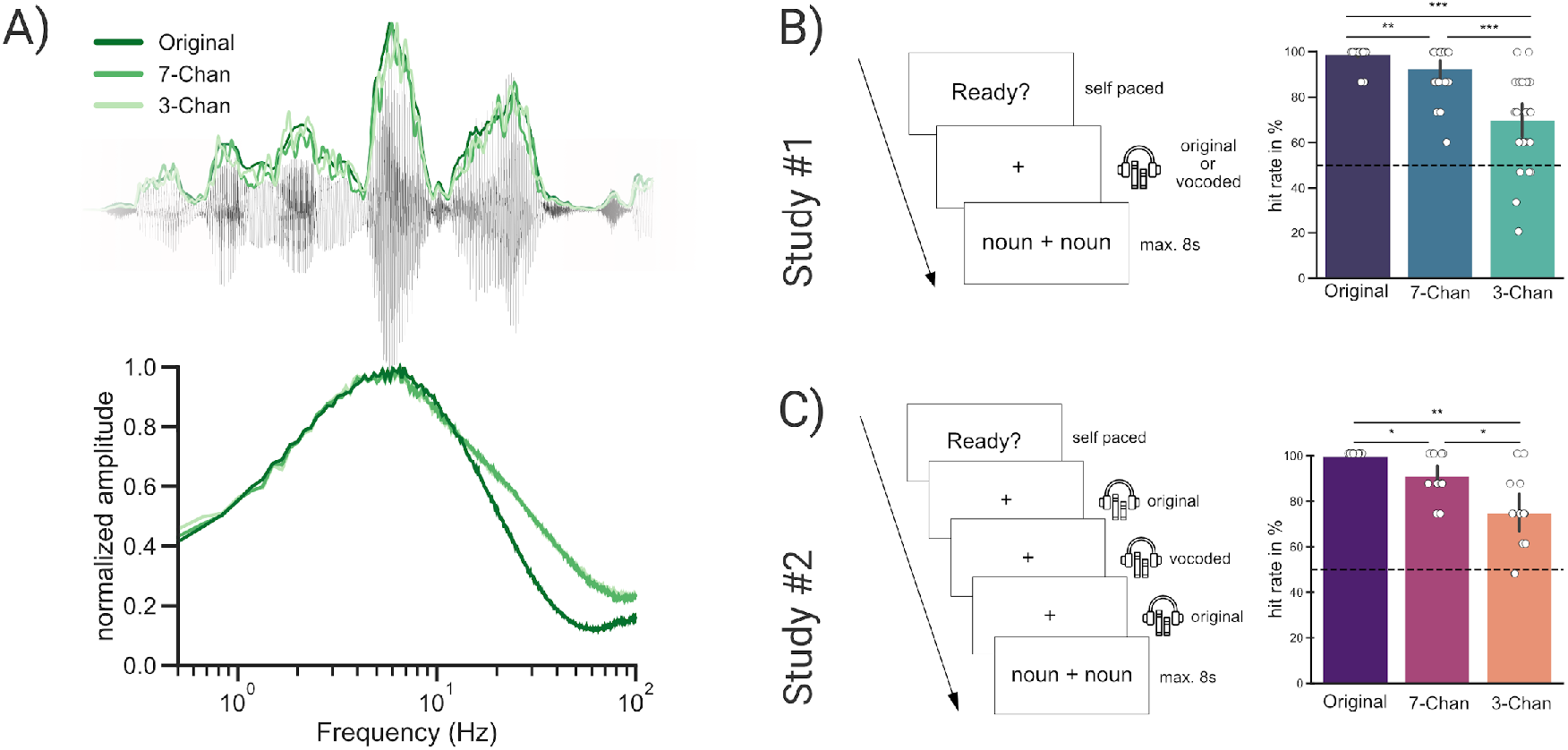
Task performance declines with speech intelligibility. (**A**) An excerpt from the audiobook presented with the corresponding speech envelope (Original) and with the envelopes of the vocoded audio stimuli (7-Chan, 3-Chan) and the averaged modulation spectra of the audio streams. (**B**) In Study#1 subjects listened either to a continuous segment of clear or vocoded speech. (**C**) In Study#2 short segments of vocoded speech ∼6-18 s were embedded in an otherwise clear speech stream (∼1-3 minute duration). In both studies, subjects were presented with two nouns at the end of each stimulus. They were further instructed to pick the one they perceived in the previous sentence. Hit rates declined in both experiments with a decrease in speech intelligibility.

Task performance declined in both experiments with speech intelligibility, recognizable by a decrease in the mean hit rate. A one-way repeated measures ANOVA across the three conditions revealed a primary effect for Study#1 (*F*(2, 48) = 44.583, *p*_*ggeisser*_ = 7.35e^-09^, *η*_*p*_^*2*^ = 0.65) and Study#2 (*F*(2, 26) = 24.536, *p* = 1e^-06^, *η*_*p*_^*2*^ = 0.654). Comparing the different vocoding levels with each other showed higher hit rates for unvocoded stimuli than for stimuli vocoded with 7-Channels (Study#1, *z*(24) = 2.916, *p*_*fdr*_ = 0.0035, *d* = 0.853; Study#2, *z*(13) = 2.566, *p*_*fdr*_ = 0.0102, *d* = 1.39) or 3-Channels (Study#1, *z*(24) = 3.955, *p*_*fdr*_ = 7.7e^-05^, *d* = 2.151; Study#2, *z*(13) = 2.720, *p*_*fdr*_ = 0.0065, *d* = 2.280). Whereas stimuli vocoded with 7-Channels showed higher hit rates than stimuli vocoded with 3-Channels (Study#1, *z*(24) = 3.955, *p*_*fdr*_ = 0.0002, *d* = 1.491; Study#2, *z*(13) = 2.572, *p*_*fdr*_ = 0.0101, *d* = 1.265). Across all conditions hit rates differed significantly from chance (Study#1, Fig. 1B; Study#2, Fig. 1C): for unvocoded speech (Study#1, *z*(24) = 4.838, *p*_*fdr*_ = 3.932e^-06^; Study#2, *z*(13) = 3.742, *p*_*fdr*_ = 0.0005), for 7 vocoding channels (Study#1, *z*(24) = 4.483, *p*_*fdr*_ = 1.103e^-05^; Study#2, *z*(13) = 3.355, *p*_*fdr*_ = 0.0011) and for 3 vocoding channels (Study#1, *z*(24) = 3.625, *p*_*fdr*_ = 0.0003; Study#2, *z*(13) = 3.105, *p*_*fdr*_ = 0.0019). This shows that while speech comprehension gradually decreases with increases in vocoding, speech was still intelligible even when only 3-Channels were used to vocode the presented audio files.

### Speech brain coherence declines with speech intelligibility

To investigate how a loss of speech intelligibility via noise-vocoding influences the neural dynamics of speech tracking we measured the coherence between the speech envelope and the related cortical activity (see coherence spectra in Fig. 2A). Comparisons of the coherence spectra across the three conditions (Original, 7-Channels and 3-Channels) using a cluster-corrected repeated-measures ANOVA, revealed a significant difference in the low-frequency range (averaged between 2 and 7Hz) for both Study#1 (*p* = 0.0004) and Study#2 (*p* = 9e^-05^). This difference was strongest in right superior temporal gyrus for both Study#1 and #2. Both in Study#1 and #2 listening to the unaltered audio resulted in the strongest speech-brain coherence, while the stimuli with a strong degradation (3-Channels) elicited the weakest coherence (Fig. 2C & 2F). Listening to the unaltered (“Original”) audio files elicited stronger speech-brain coherence than listening to speech vocoded with 7-Channels in Study#1 (*t*(27) = 2.519, *p*_*fdr*_ = 0.018, *d* = 0.467) but not in Study#2 (*t*(26) = 1.425, *p*_*fdr*_ = 0.166, *d* = 0.307). However, listening to the unaltered (“Original”) audio files elicited a stronger coherence than listening to speech in the 3-Channel condition (Study#1, *t*(27) = 6.083, *p*_*fdr*_ = 3e^-06^, *d* = 1.623; Study#2, *t*(26) = 7.451, *p*_*fdr*_ = 1.959e^-07^, *d* = 1.787). Listening to the 7-Channels condition elicited higher levels of speech-brain coherence than listening to the 3-Channels condition (Study#1, *t*(27) = 6.238, *p*_*fdr*_ = 3e^-06^, *d* = 1.446; Study#2, *t*(26) = 7.021, *p*_*fdr*_ = 2.802e^-07^, *d* = 1.599).

**Fig 2.**
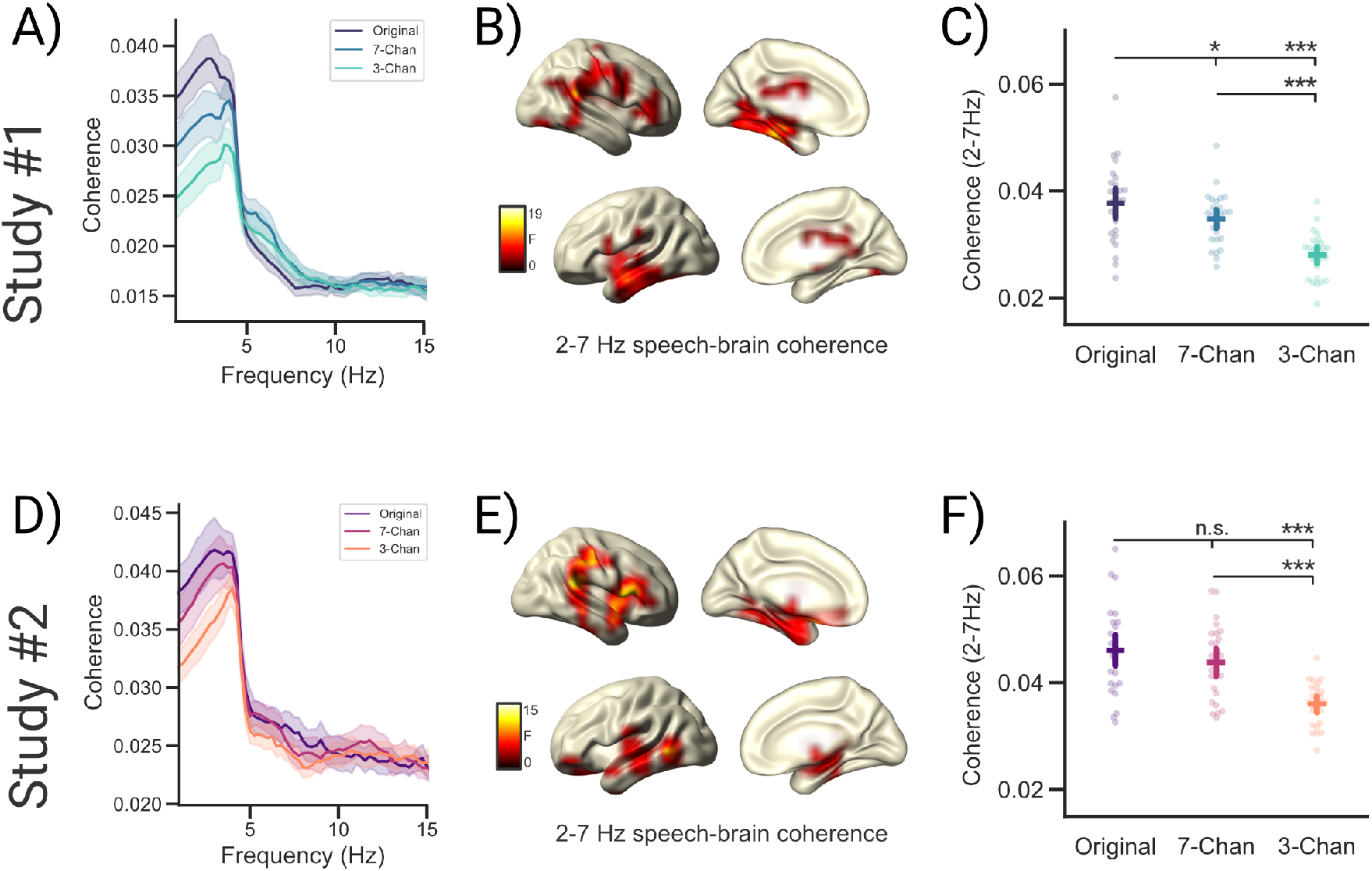
Speech brain coherence declines with speech intelligibility. (**A**,**D**) Speech-brain coherence spectra for the three conditions averaged across all voxels. (**B**,**E**) Source localizations of degradation effects on speech-brain coherence (2-7Hz) during acoustic stimulation across three conditions (Original, 7-Chan & 3-Chan) in bilateral temporal and medial frontal regions. (**C**,**F**) Individual coherence estimates (averaged) of the three vocoding conditions extracted at voxels showing a significant difference using a cluster-corrected permutation test. Bars represent 95% confidence intervals, p_fdr_ < 0.05*, p_fdr_ < 0.01**, p_fdr_ < 0.001***

In sum, these results show that both intermittent and continuous degradation similarly affect low-frequency speech brain coherence. In both experimental designs, speech brain coherence decreased as speech became less intelligible. Comparing the decrease in coherence through vocoding across studies revealed that this decrease was not different across both studies (*U* = 297, *p* = .175, *r* = .214). At first glance, these results are in conflict with a previous analysis of Study#1 (*11*). The main difference between the previous and the current analysis of Study#1 can primarily be attributed to different filter settings (lower cut-off for the high-pass filter in the current analysis) during preprocessing that affected the offset and exponent of the speech-brain coherence spectrum differently (see Discussion). In the present study, these changes were applied to allow for better modeling of the periodic and the aperiodic components of the coherence spectrum. Interestingly, further analysis of these components showed that the aperiodic components explain most of the variance (Offset/Exponent; Study#1, *r*^*2*^ = 0.83/0.67; Study#2, *r*^*2*^ = 0.36/0.32) of the averaged (2-7Hz) low-frequency speech-brain coherence in both studies (see Supplementary Materials; Fig. S2). This illustrates that analysing coherence differences in a band-limited range may be strongly influenced by aperiodic differences that do not necessarily reflect neural tracking of sound or linguistic information in the relevant frequency range. Depending on the filter settings, these aperiodic components may heavily impact the results. This observation is especially important for investigations that focus on slow and infraslow modulations and highlights the necessity to separate periodic from aperiodic contributions.

### Declining speech intelligibility increases the center frequency of neural speech tracking along with a sharper tuning

Both the speech envelope and electrophysiological signals (recorded using EEG/MEG) are characterized by an overall 1/f-like spectrum (*13, 14*). This appears to also be evident in the coherence estimation between both signals (independent of the speech-relevant peak at low frequencies; see Fig. 2A,D). To quantify relevant aspects of the periodic components of speech tracking we extracted the most prominent peaks of the coherence spectra in the low-frequency range across all virtual channels in which we observed a significant difference across vocoding levels (see Fig. 2B, E). This was operationalized by using FOOOF (*12*) to first flatten the coherence spectrum and then compute Gaussian model fits to extract peaks. For each subject, the average relative magnitude of the coherence peak, the bandwidth (∼tuning) and center frequency of the extracted peaks (Fig. 4) were computed and compared within subjects and across the three conditions (Original, 7-Channels and 3-Channels) using a repeated-measure ANOVA.

**Fig 3.**
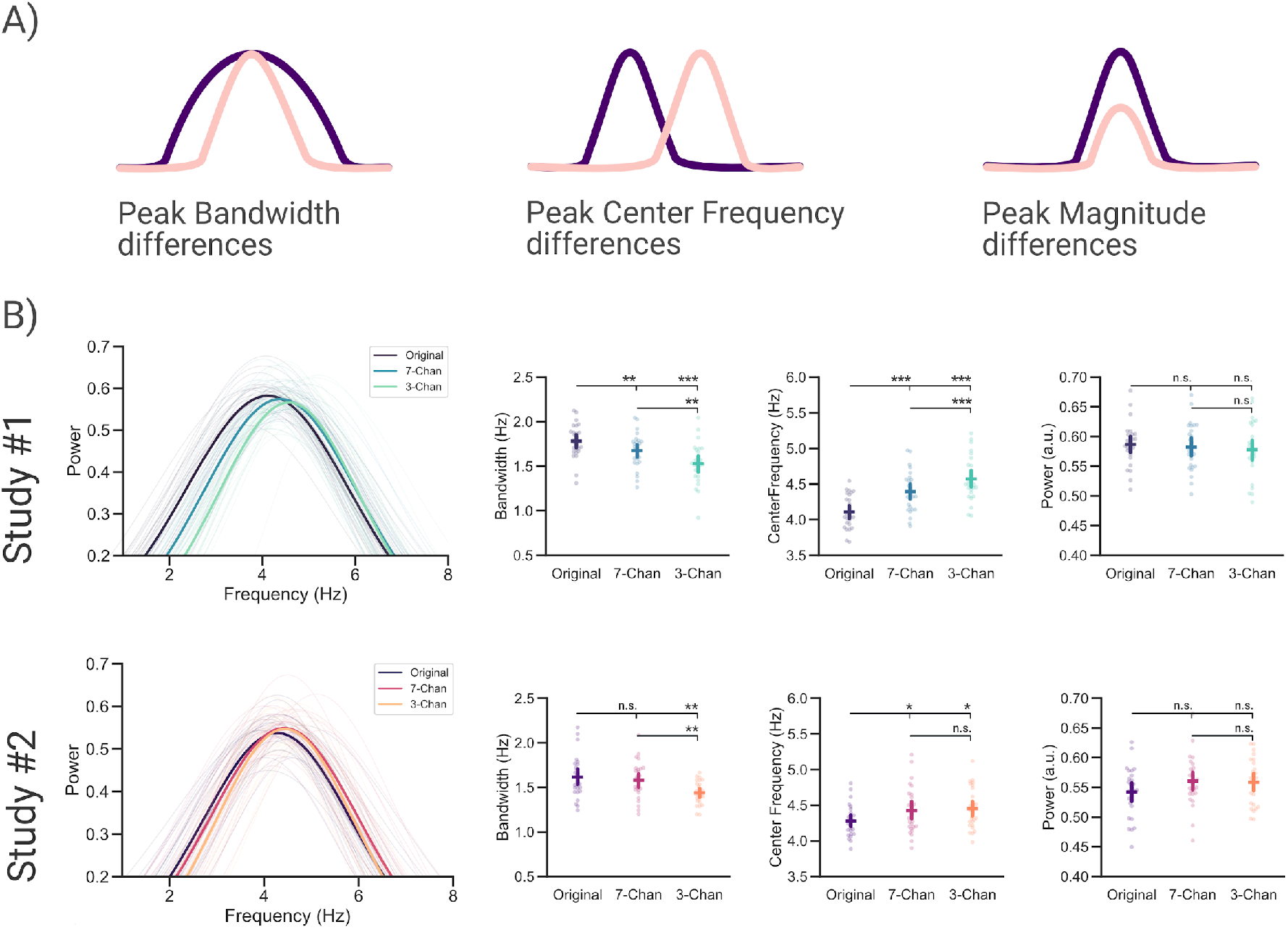
Declining speech intelligibility increases the center frequency of neural speech tracking along with a sharper tuning. (**A**) Peak parameters influencing a significant coherence difference across experimental conditions. (**B**) The averaged relative magnitude, center frequencies and bandwidth of peaks extracted from the coherence spectra for each subject were compared across three conditions (Original, 7-Chan & 3-Chan). Bars represent 95% confidence intervals, p_fdr_ < 0.05*, p_fdr_ < 0.01**, p_fdr_ < 0.001***

**Fig 4.**
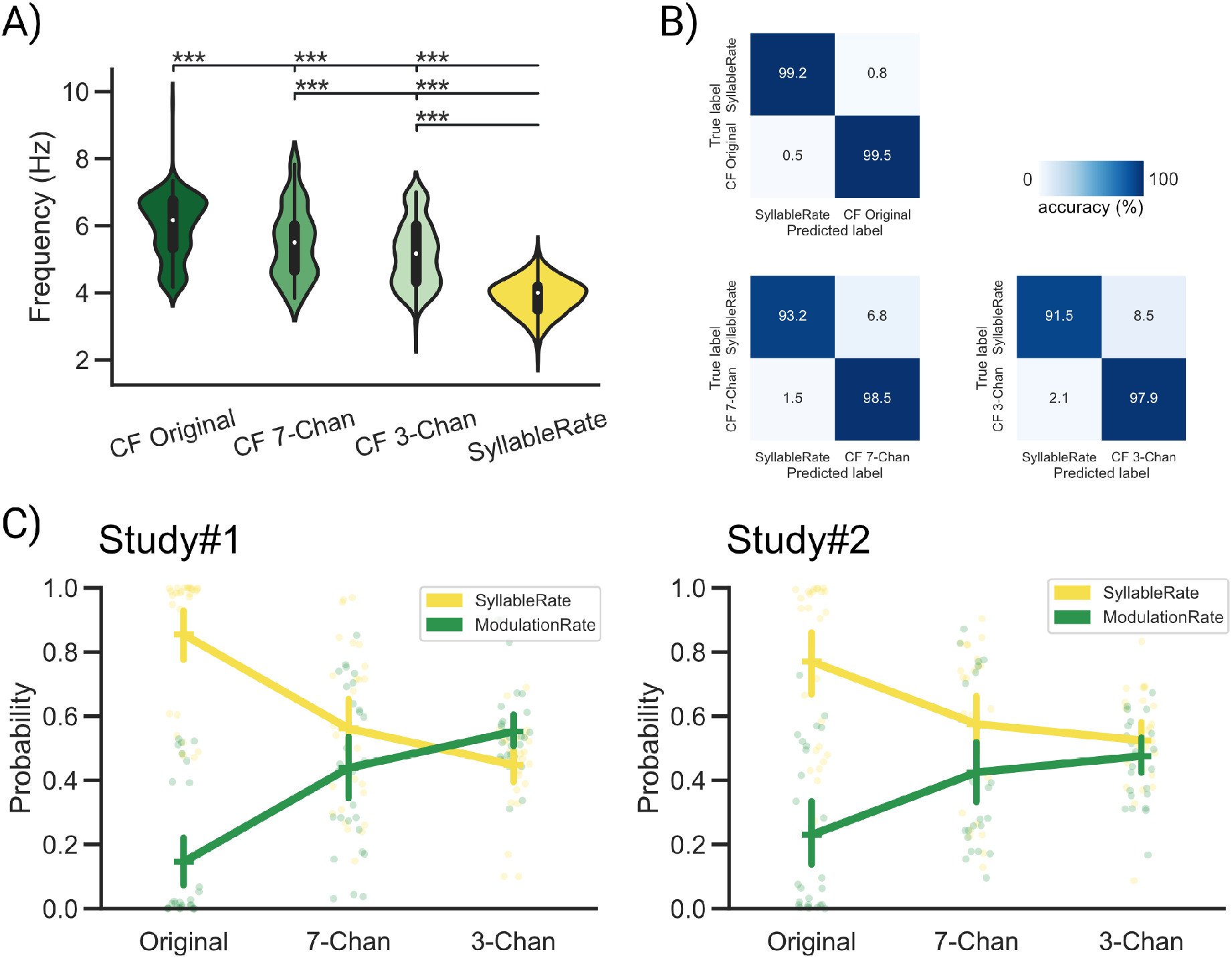
Neural speech tracking shifts from syllabic to modulation rate as speech intelligibility decreases. (**A**) Center frequencies extracted from the acoustic envelopes of clear and vocoded speech (green) and the syllabic rate (yellow). Although, we generally noted a strong overlap across the modulation spectra (see Fig. 1A) the extracted center frequencies of the acoustic envelopes differed not only significantly from the syllabic rate, but also across vocoding levels. (**B**) An ensemble of k-nearest neighbors classifiers were trained in a nested cross validation scheme to decode the modulation rate vs. the syllabic rate. (**C**) Classifiers applied to the center frequencies of the coherence spectra to decode whether the tracking was either related to the tracking of the syllable or the modulation rate of speech. Bars represent 95% confidence intervals, p_fdr_ < 0.05*, p_fdr_ < 0.01**, p_fdr_ < 0.001***

This analysis showed that the actual magnitude of the extracted peaks did not differ across the three vocoding conditions in both studies (Study#1, *F*(2, 54) = 0.522, *p* = 0.596, *η*_*p*_^*2*^ = 0.019; Study#2, *F*(2, 50) = 2.18, *p* = 0.124, *η*_*p*_^*2*^ = 0.08). However, we noticed a significant difference across the center frequencies of the detected peaks over the three conditions in both studies (Study#1, *F*(2, 54) = 48.628, *p* = 8.365e^-13^, *η*_*p*_^*2*^ = 0.643; Study#2, *F*(2, 50) = 5.28, *p* = 0.008, *η*_*p*_^*2*^ = 0.175). Comparing the different vocoding levels with each other showed lower center frequencies for unvocoded stimuli than for stimuli vocoded with 7-Channels (Study#1, *t*(27) = −7.122, *p*_*fdr*_ = 1.753e^-07^, *d* = −1.271; Study#2, *t*(25) = −2.756, *p*_*fdr*_= 0.016, *d* = −0.613) and with 3-Channels (Study#1, *t*(27) = −8.797, *p*_*fdr*_ = 6.18e^-09^, *d* = −1.918; Study#2, *t*(25) = −2.946, *p*_*fdr*_ = 0.0161, *d* = −0.7). The two vocoding conditions did differ significantly from each other in Study#1 (*t*(27) = −3.227, *p*_*fdr*_ = 3.273e^-03^, *d* = −0.544) but not in Study#2 (*t*(25) = −0.114, *p*_*fdr*_ = 0.91, *d* = −0.023) with lower center frequencies for speech vocoded with 7-Channels compared to speech vocoded with 3-Channels.

For the bandwidth of the detected peaks, differences across the three conditions were also observed both in Study#1 (*F*(2, 54) = 18.808, *p* = 6.329e^-07^, *η*_*p*_^*2*^ = 0.411) and Study#2 (*F*(2, 50) = 5.444, *p* = 0.007, *η*_*p*_^*2*^ = 0.179). In the continuous design, (Study#1) the tuning bandwidth for unvocoded stimuli was broader than for stimuli vocoded with 7-Channels (*t*(27) = 3.219, *p*_*fdr*_ = 0.003, *d* = 0.666) and with 3-Channels (*t*(27) = 5.196, *p*_*fdr*_ = 5.4e^-05^, *d* =1.422). In the intermittent design (Study#2), the direction of the effect was similar, yet only significant for the difference between unvocoded speech and speech vocoded with 3-Channels (*t*(25) = 3.398, *p*_*fdr*_ = 0.007, *d* = 0.983) and not for the difference between unvocoded speech and speech vocoded with 7-Channels (*t*(25) = 0.699, *p*_*fdr*_ = 0.491, *d* = 0.201). Speech vocoded with 7-Channels had a broader tuning bandwidth than speech vocoded with 3-Channels across both studies (Study#1, *t*(27) = 3.592, *p*_*fdr*_ = 0.002, *d* = 0.758; Study#2, *t*(25) = 2.668, *p*_*fdr*_ = 0.02, *d* = 0.774).

In sum, these results show that intermittent and continuous degradation similarly affect the periodic components of speech-brain coherence that are putatively reflective of neural speech tracking. Interestingly, the difference between speech tracking across different levels of intelligibility was not driven by the relative height of the peak in the coherence spectrum, but rather by a sharper tuning (Fig. 4; Bandwidth) combined with an increase of center frequencies of the coherence spectra (Fig. 4; Center Frequency).

### Neural speech tracking shifts from syllabic to modulation rate as speech intelligibility decreases

As speech intelligibility decreases we noted an increase of the center frequencies of speech-brain coherence. We also extracted the center frequencies of the modulation spectra from the acoustic envelopes of the audiobook for the three conditions (Original, 7-Channels, 3-Channels as in (*9*); see Fig. 4A) and computed the realized syllable rate of the presented audiobook (*18*). Although, there was generally a strong overlap over the modulation spectra of the speaker across vocoding levels (see Fig. 1A), a one-way repeated measures ANOVA across the extracted center frequencies and the syllable rate of the audio signal revealed a significant main effect (*F*(3, 1098) = 454.104, *p* = 2.68e^-175^, *η*_*p*_^*2*^ = 0.554). All conditions differed significantly from each other (see Fig. 4 & Table S1 for a related post-hoc analysis). The rate at which the syllables were produced (*Mdn* = 4 Hz) was lower than the center frequencies of the modulation spectra of the audio signal 3-Channels (*Mdn* = 5.16 Hz), 7-Channels (*Mdn* = 5.5 Hz) and clear speech condition (*Mdn* = 6.16 Hz). The increase in center frequencies of speech-brain coherence along with the differences in modulation and syllable rates suggests that the brain may be driven more by acoustic or linguistic information depending on the signal quality. This is intuitive, as with increased vocoding it also becomes more difficult to extract linguistically meaningful information such as phrase boundaries or syllables. This mainly leaves the modulation intensities of the acoustic speech envelope as an information source to the listener. The following analysis aims at addressing this point more directly.

We trained and tested an ensemble of *k*-nearest neighbor classifiers to test whether neural speech tracking shifts from the syllabic (linguistic information) to the modulation rate (acoustic information) as speech becomes less intelligible. This analysis was performed in a nested 5-fold cross-validation (see Methods for a detailed account) to differentiate between the center frequencies of the modulation spectra for the three conditions (Original, 7-Channels, 3-Channels) and the realized syllable rate of the speaker (see Fig. 4A). The results of the nested cross-validation procedure (Fig. 4B) show that the classifiers can predict with a high accuracy whether a given frequency in hertz can be related either to the modulation or realized syllable rate of the speaker. We then used the weights of these classifiers to predict whether the extracted center frequencies of speech brain coherence were related more closely to the realized syllable rate or the modulation rate of our speaker. This analysis showed that in the unaltered clear speech condition, neural speech tracking was closely related to the syllable rate. However, as intelligibility decreases the probability that the classifiers predict that a given center frequency is related rather to the modulation as opposed to the syllabic rate increases.

The results of a two-way repeated measures ANOVA revealed that there was a significant main effect for the factors tracking (Modulation/Syllable rate) in both studies (Study#1 *F*(1, 27) = 16.175, *p*_*ggeisser*_ = 0.0004, *η*_*p*_^*2*^ = 0.375; Study#2 *F*(1, 25) = 18.999, *p*_*ggeisser*_ = 0.0002, *η*_*p*_^*2*^ = 0.432). The probability that neural speech tracking is reflective of the syllable rate (linguistic component) was overall higher than the tracking of the modulation rate (acoustic component). There was no significant main effect of Vocoding (Original, 7-Channels, 3-Channels; Study#1 *F*(2, 54) = 0, *p* = 1; Study#2 *F*(2, 48) = 0, *p* = 1) this is intuitive as the overall probability in each condition is 0.5 when ignoring the factor tracking (Modulation/Syllable rate). However, there was a significant interaction effect for the factors tracking (Modulation/Syllable rate) and vocoding (Original, 7-Channels, 3-Channels) across both Studies (Study#1 *F*(2, 54) = 47.340, *p*_*ggeisser*_ = 1.387e^-11^, *η*_*p*_^*2*^ = 0.637; Study#2 *F*(2, 50) = 10.235, *p*_*ggeisser*_ = 0.0006, *η*_*p*_^*2*^ = 0.29). This suggests that while speech intelligibility decreases and less linguistically meaningful information is present, neural speech tracking starts to drift away from the syllabic rate towards the modulation rate of speech.

### Modelling of subcortical activity reveals a predominant tracking of the modulation rate of speech

Recent studies using non-invasive electrophysiology have shown that auditory activity at putative subcortical processing stages can be measured for complex natural sounds (such as speech; *19*–*22*). Furthermore, this subcortical activity can even be modulated by attention (*19, 20, 23*). Interestingly, top-down attentional modulations of auditory activity can already be detected at the hair cells in the inner ear measured as otoacoustic activity (faint sounds emitted by the outer hair cells; see *24*). Other studies have shown that even subcortical nuclei on the auditory pathway are behaviorally relevant for speech recognition (medial geniculate bodies; *25*). Using a recently developed modeling procedure (*17*), we further aimed to investigate whether differences in speech intelligibility can be already observed at putative subcortical processing stages.

We used a localizer measurement (*17*) to compute individualized weights (per subject; note that the localizer was only available for Study#2). These weights reflect activity along the auditory hierarchy, resulting in 100 virtual channels ranging putatively from the auditory nerve (channels 0-20) to early thalamo(-cortical) processing stages (channels 90-100). We then applied these weights (see Material & Methods: Modelling of subcortical auditory activity) to the epoched data from Study#2 to infer activity along the auditory hierarchy (see spectral distribution in Fig. 5A). A cluster-corrected repeated-measures ANOVA across the three conditions (Original, 7-Channels and 3-Channels) and within subjects revealed a significant difference in the low frequency range (2-7Hz) between virtual channels that are reflective of subcortical activity at early stages of auditory processing (putatively auditory nerve/cochlear nucleus, *p* = 0.0045). Listening to the the unaltered (“Original”) audio files elicited higher speech-brain coherence than listening to the 7-Channels (*t*(24) = 3.2, *p*_*fdr*_ = 0.005, *d* = 0.798) and the 3-Channels condition (*t*(24) = 4.282, *p*_*fdr*_ = 0.0008, *d* = 1.212). However, the two vocoding conditions did not differ significantly from each other (*t*(24) = 1.547, *p*_*fdr*_ = 0.135, *d* = 0.488).

**Fig 5:**
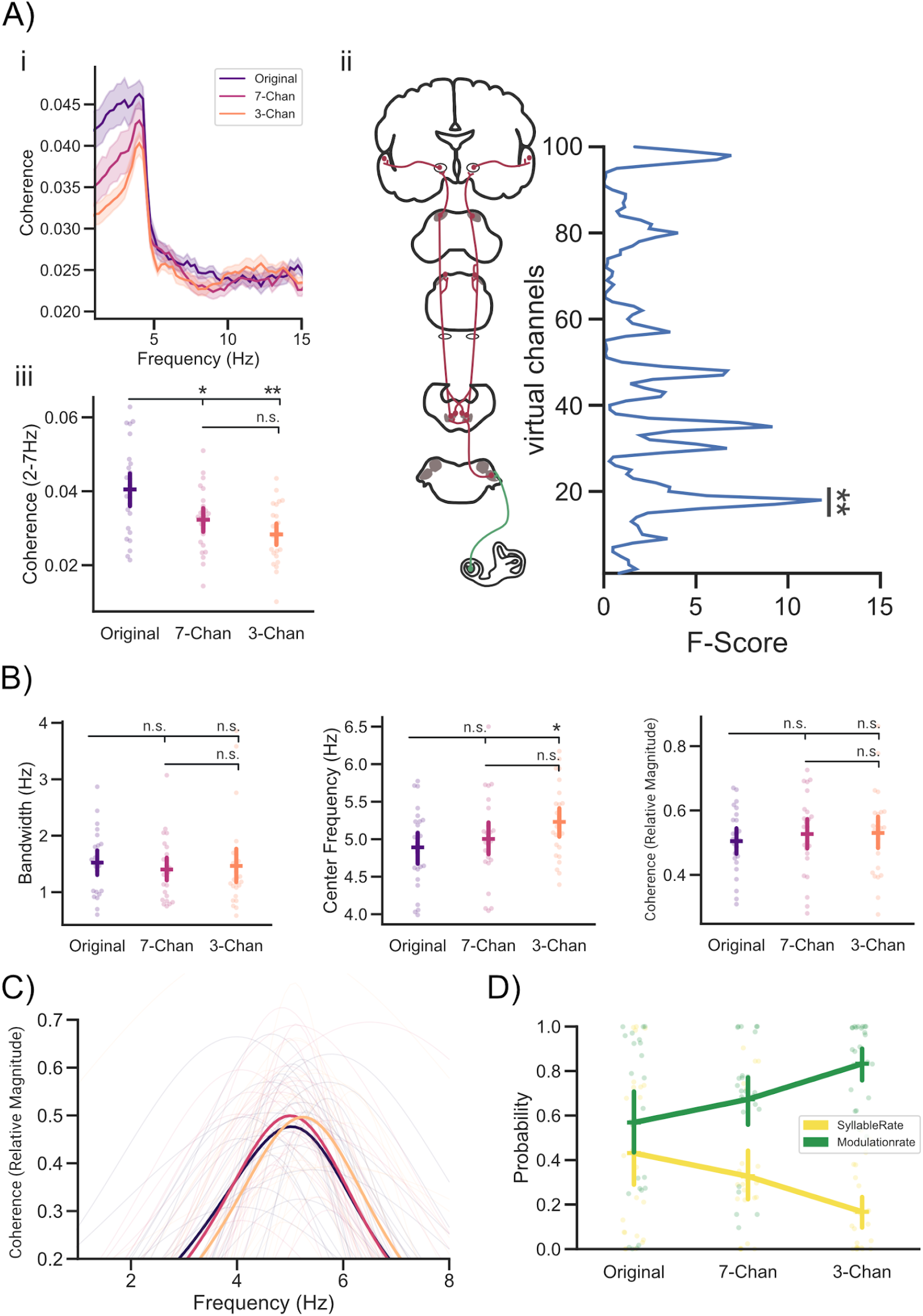
Modelling of subcortical activity reveals a predominant tracking of the modulation spectrum of speech. (**Ai**) Speech-brain coherence spectra for the three conditions averaged across all virtual channels. (**Aii**) The three conditions (Original, 7-Chan & 3-Chan) differed significantly at virtual channels reflecting activity putatively related to the auditory nerve/cochlear nucleus (channels 16-20). (**Aiii**) Individual coherence (2-7Hz) extracted at channels 16-20 was highest for clear speech. (**B**,**C**) Peak height, center frequencies and bandwidth of peaks extracted from virtual channels (16-20) (**D**) Classifiers applied to the center frequencies of the coherence spectra to decode whether speech tracking was either related to the tracking of the syllable or the modulation rate of speech. Bars represent 95% confidence intervals, p_fdr_ < 0.05*, p_fdr_ < 0.01**, p_fdr_ < 0.001***

We further investigated the periodic components that are reflective of speech tracking by extracting peaks from the coherence spectra to analyse the corresponding magnitude of the coherence peak, the bandwidth and center frequencies. A repeated-measures ANOVA across conditions (Original, 7-Channels and 3-Channels) and within subjects revealed no significant differences for the relative magnitude of the coherence peak (*F*(2, 48) = 0.335, *p* = 0.717, *η*_*p*_^*2*^ = 0.014) and the bandwidth of the extracted peaks (*F*(2, 48) = 0.192, *p* = 0.826, *η*_*p*_^*2*^ = 0.008). However, significant differences were found across conditions for the center frequencies of the peaks (*F*(2, 48) = 3.213, *p* = 0.049, *η*_*p*_^*2*^ = 0.118). Listening to the unaltered (“Original”) audio files was associated with significantly lower center frequencies than listening to the 3-Channel condition (*t*(24) = −3.062, *p*_*fdr*_ = 0.016, *d* = −0.664) but not than listening to the 7-Channel condition (*t*(24) = −0.767, *p*_*fdr*_ = 0.450, *d* = −0.204) at subcortical processing stages. The two vocoding conditions did not differ significantly from each other (*t*(24) = −1.528, *p*_*fdr*_ = 0.21, *d* = −0.425).

We applied the pre-trained classifiers (see Fig. 4) to detect whether the tracking of speech at putatively early auditory processing stages could be either related to the modulation rate of speech or the syllabic rate. We found that modelling of the related subcortical activity reveals predominantly a tracking of the acoustic modulation rate of speech (*F*(2, 48) = 23.220, *p* = 6.6e^-05^, *η*_*p*_^*2*^ = 0.492), contrary to the previous analysis mainly reflecting cortical effects (see Fig. 4). However, similar to the previous analysis reflecting mainly cortical activity there was an interaction effect between tracking and vocoding, with decreasing intelligibility the probability increases that the classifiers rather predict that the center frequency is related to the modulation as opposed to the syllabic rate (*F*(2, 48) = 6.947, *p* = 0.00224, *η*_*p*_^*2*^ = 0.224).

In sum, these results suggest that differences in speech tracking between clear and vocoded stimuli already arise at subcortical processing stages. This difference in neural speech tracking occurs at virtual channels that can be associated with subcortical activity between the auditory nerve and cochlear nucleus. The extracted peaks at this level of processing did not differ significantly from each other regarding the relative peak height of the coherence (similar to cortical observations; see Fig. 4) and their tuning width (different to cortical observations; see Fig. 4). Yet, the center frequency shift of these peaks showed a similar effect when compared to cortical processing stages. As intelligibility decreased the probability that the center frequencies could be related to the modulation opposed to the syllable rate increased steadily. However, contrary to the cortical recordings, tracking across different levels of intelligibility at subcortical processing stages was predominantly related to the modulation rate of speech rather than the syllabic rate. This shows that although tracking at a subcortical level is dominated by lower-level acoustic envelope modulations, intelligibility also influences these hierarchically early responses.

## DISCUSSION

Speech tracking is modulated by the intelligibility of the sensory input. However, the pattern of that modulation-frequently operationalized by band-limited coherence effects- is not consistent across studies (see e.g. *9, 11, 26*). This complicates a mechanistic understanding of how speech tracking actually supports speech comprehension. Applying a method to separate periodic from aperiodic components in the coherence spectrum, our results yield a differentiated picture, indicating that intelligibility affects tuning-width and center frequency of the periodic components in the low frequency range.

### Band-limited speech-brain coherence declines with speech intelligibility

Here, we investigated the effects degraded speech has on the neural dynamics of speech tracking using data from two slightly different experimental paradigms. In Study#1 speech was displayed continuously at one of three different levels of intelligibility (Original, 7-Channels, 3-Channels; ∼15s-3min). In Study#2 segments of degraded speech (7-Channels, 3-Channels; ∼6-18s) were embedded in a clear audio stream (Original; ∼1-3min) as both studies produced comparable results, they will be discussed together. We observed in accordance with previous results (*1*–*4*) that low frequency speech-brain coherence declines with a decrease in intelligibility. However, other studies have reported a variety of partly contradicting results (*9*–*11*). Our present results show that the reported band-limited coherence spectra are very strongly related to the underlying aperiodic components in the spectrum (see supplementary material). Since the field is mostly interested in neural tracking of (relatively) periodic speech features around the syllable rate, it is questionable whether band-limited coherences without consideration of the aperiodic components are a viable measure for neural speech tracking.

### Neural speech tracking shifts from syllabic to modulation rate as speech intelligibility decreases

Interestingly, in the investigation of spectral power differences in electrophysiological signals, a variety of contradicting results is also commonly reported for band-limited effects. This appears to be caused by the conflation of periodic (center frequency, power, bandwidth) and aperiodic (offset, exponent) properties of the underlying signal (*12*). This is deemed problematic as periodic and aperiodic components of the signal can be linked to a variety of different effects (*12*). Both the acoustic envelope of speech and electrophysiological measurements of neural activity possess an overall (aperiodic) 1/f-like spectrum (*13, 14*). This 1/f-like pattern is at times also found in the low-frequency coherence/correlation spectrum between both signals (e.g. see *1, 9, 11*). We therefore decomposed the speech-brain coherence spectra in their periodic and aperiodic components using FOOOF (*12*), to better understand the relationship between the intelligibility of speech and the related neural dynamics of speech tracking. Interestingly, these investigations revealed that the aperiodic components (offset & exponent) explained most of the variance observed in the coherence difference (at 2-7 Hz; see Fig. 2) across vocoding levels (see Fig. S2). This highlights the importance of separating periodic from aperiodic components in the speech-brain coherence spectra, as we were primarily interested in investigating peaks in the coherence spectra (periodic components) that can be related to neural speech tracking. Further investigations of the periodic components of the low frequency coherence peak (center frequency, relative magnitude, bandwidth) revealed that there was no difference across vocoding levels in the relative magnitude of the coherence peak. Instead, the differences in neural speech tracking were rather caused by a sharpening in the frequency tuning of the coherence peak of vocoded speech along with an increase of the center frequencies of the observed peaks (see Fig. 3). Using a decoding analysis, we were able to link the increase of the center frequencies to a shift in tracking from higher-level linguistic to lower-level acoustic information of the speech stream. Our analysis showed that as intelligibility decreases the probability that tracking is related to the modulation (acoustic) rather than the syllabic rate (linguistic) of speech increases. This is intuitive, as with decreased intelligibility it also becomes more difficult to extract linguistically meaningful information such as phrase boundaries or syllables. This mainly leaves the modulation intensities of the acoustic speech envelope as an information source to the listener. As the acoustic modulation of speech is closely related to the production of syllables (*5*), investigations of neural speech tracking are typically not making a distinction between lower-level acoustic and higher-level linguistic information on the level of syllable processing. However, while the modulation rate (acoustic property) of speech appears to be exceptionally stable across languages and speaking conditions (*5, 6*), the syllable rate (linguistic property) of speech differs depending on the language and the speaking conditions (*7, 8*). This suggests that modulation rate and syllable rate are not terms that can be necessarily used interchangeably. Therefore, distinguishing these properties more clearly may be important to gain a better understanding of the neural processes separating auditory processing disorders (e.g. hearing loss) from language processing disorders (e.g. developmental dyslexia), which has been difficult based solely on neural speech tracking. This difficulty may be linked to the variety of (partly contradicting) results within and across auditory/linguistic processing disorders that relate to the neural dynamics of speech tracking. While a recent study was able to link hearing loss to a relative increase in speech envelope tracking (compared to (age matched) normal hearing listeners (*27*)), previous studies could not report enhanced envelope tracking in individuals with a hearing-impairment (*28, 29*). Related to language proficiency, similar inconsistencies are reported as non-native speakers appear to show an increased envelope tracking compared to native speakers (*10, 30*). On the other hand, individuals suffering from developmental dyslexia are reported to have lower synchronization with the speech envelope compared with neurotypical individuals (*31*). This range of (partly contradicting) results again highlights the complex relationship between the intelligibility of speech and the related neural dynamics of speech tracking. Using the approach proposed here of decomposing coherence spectra in their periodic and aperiodic components, it should be possible to gain a more fine-grained view on the specific characteristics underlying the neural dynamics of speech tracking. This may help in the future to better differentiate the neural signatures of individuals suffering from auditory processing or language processing disorders.

### Declining speech intelligibility goes along with a sharper frequency tuning

Apart from the intelligibility-dependent changes in the center frequencies of the coherence peaks, we also noted a wider frequency tuning of speech tracking in clear as opposed to vocoded speech. The width of this frequency tuning decreased with a loss in intelligibility. As the syllabic rate of our speaker (∼4Hz) differed from the modulation rate of her speech stream (∼5-6Hz; see Fig. 4), the narrowing in tuning may also be related to a loss in linguistically meaningful information. This might suggest that in situations where speech is clear, both linguistic (syllable rate) and acoustic information (modulation rate) were tracked resulting in an increased bandwidth covering all relevant frequencies. As speech becomes less intelligible and it becomes harder to extract linguistically meaningful information, the bandwidth of the coherence peak narrows around the higher frequencies of the residual acoustic modulation of speech. Furthermore, previous studies have shown that auditory selective attention effects may arise from an enhanced tuning of receptive fields of task-relevant neural populations (*32, 33*). Therefore, the observed narrower frequency tuning could also be related to enhanced top-down auditory attention processes (*34*) in situations where listening becomes more challenging.

### Influences of aperiodic components on the neural dynamics of speech tracking

Investigating the parameters related to low frequency peaks in measurements of speech brain coherence is offering a new and unique perspective to better understand the neural dynamics underlying speech tracking. However, we also noticed that band-limited differences in the speech-brain coherence spectra are strongly related to the underlying aperiodic components. This highlights the importance to separate periodic from aperiodic components, as periodic and aperiodic components can be linked to a variety of different effects (*12*). Commonly, aperiodic components of most signals have been considered as noise and as such are often just removed from the overall signal. Especially for low-frequency activity this can be easily achieved by spectrally normalising (whitening) the signal via filtering (e.g. see *35*). Different choices in filter settings however can also generally accentuate different properties of a signal. For instance, in this study we reanalysed data from a recent study (Study#1 (*11*)) using a larger time window for the coherence estimation (4s instead of 2s; to obtain a better frequency resolution for low frequency speech tracking) and a lower cut-off for the high-pass filter (0.1Hz instead of 1Hz). These changes were intended to improve the model fit of FOOOF for the low frequency coherence spectra, but also resulted in a different pattern for low frequency speech-brain coherence (compare Fig. 2ABC) with Fig. 2ab) in (*11*)). The previous analysis of Study#1 (*11*) showed that neural speech tracking increases for mild decreases in intelligibility (putatively driven by an increased listening effort) and then decreases as speech becomes increasingly unintelligible. We now show that low frequency speech tracking gradually decreases with intelligibility. This difference was mainly driven by changes in filter settings accentuating different properties of the signal by putatively differently influencing the 1/f-like pattern of low frequency speech-brain coherence. Similar to the analysis of power spectral densities, 1/f-like patterns in the coherence spectra also appear to play a striking role when computing statistics across experimental conditions (see S2 for a comparison of slope and offset for the data analysed in the present study). However, whether or not 1/f-like patterns carry (in general) meaningful information is heavily debated. Nevertheless, recent studies have shown that 1/f-like patterns in electrophysiological power spectra can change both dependent on trait-like factors (age (*36*), ADHD (*37*) and schizophrenia (*38*)) and state-like factors (e.g. differences over cognitive and perceptual states (*39, 40*).This suggests a physiologically meaningful underpinning of 1/f-like neural activity. However, interpretations related to the aperiodic patterns found in low frequency speech-brain coherence go beyond the scope of the present study, as we were mainly focused on the distinction between the processing of the syllabic rate and the modulation rate of speech related to peaks in the speech-brain coherence spectra (periodic components). Perhaps aperiodic components of speech-brain coherence could be modulated by slower components in the speech stream reflecting higher level information (e.g. sentence or phrasal information), that become increasingly lost with less intelligibility. Addressing this question should be the topic of future investigations using paradigms in which these features are parametrically controlled (*41*). However, the present study illustrates that analysing coherence in a band-limited range, even though more or less explicitly assumed, may not reflect neural tracking of sound or linguistic information in the relevant frequency range. Instead, depending on the filter settings, the aperiodic components may heavily impact the results. This is especially important for investigations that focus on slow and infraslow modulations.

### Modelling of subcortical activity reveals a predominant tracking of the modulation spectrum of speech

Previous research has shown that not only cortical, but also subcortical regions play an important role in language processing (*42*). These subcortical regions appear to be even behaviorally relevant for speech recognition (medial geniculate bodies; *25*). Here, we generated individualized spatial filters reflective of subcortical auditory processing using a localizer measurement (*17*). In principal, these filters can be applied to a separate measurement to infer subcortical auditory activity. Using this modeling procedure, we aimed to investigate whether differences in speech intelligibility can already be observed at putative subcortical processing stages. Similarly to the activity from cortical processing stages, we noticed a shift of the center frequency of the extracted peaks. As intelligibility decreased, the center frequencies of the detected peaks increased steadily. However, contrary to the cortical recordings, the applied decoding analysis showed that the center frequencies of the speech-brain coherence peaks (reflecting neural speech tracking) across different levels of intelligibility at subcortical processing stages was predominantly related to the modulation rate of speech opposed to the syllabic rate. This shows that although tracking at a subcortical level is overall higher for the low-level acoustic envelope modulation, intelligibility also influences these hierarchically early responses (see Fig. 5D). This highlights the potentially important yet often overlooked role of subcortical nuclei in speech and language processing.

## CONCLUSION

In this study, we introduce a novel way to investigate neural speech tracking by utilizing an approach recently introduced to parametrize electrophysiological power spectra (*12*). Our results show that cortical regions mostly track the syllable rate, whereas subcortical regions are driven by the acoustic modulation rate. Furthermore, the less intelligible speech becomes, the more dominant the tracking of the modulation rate becomes. Our study underlines the importance of making a distinction between the acoustic modulation and syllable rate of speech and provides novel possibilities to better understand differences between auditory processing and speech/language processing disorders. In general, parametrization of coherence spectra may offer a new and unique perspective to investigate the parameters that drive neural speech tracking across a variety of listening situations.

## MATERIALS & METHODS

### Subjects

Twenty-eight individuals participated in Study#1 (female = 17, male = 11). Mean age was 23.82 years (standard deviation, *SD* = 3.71) with a range between 19 and 37 years. In Study#2 twenty-seven individuals participated (female = 11, male = 16). Due to technical difficulties one subject was removed from Study#2. Mean age was 23.38 years (*SD* = 4.15) with a range between 19 and 38 years. Across both studies we recruited only German native speakers and people who were suitable for MEG recordings, that is, without nonremovable ferromagnetic metals in or close to the body. Participants provided informed consent and were compensated monetarily or via course credit. Participation was voluntary and in line with the declaration of Helsinki and the statutes of the University of Salzburg. The study was approved by the ethical committee of the University of Salzburg.

### Stimuli

For the MEG recording, audio files were extracted from audio–visual recordings of a female speaker reading Goethe’s “Das Märchen” (“The Tale”; 1795). In Study#1, lengths of 12 stimuli varied between approximately 15 s and 3 min, with two stimuli of 15, 30, 60, 90, 120 and 150 s, and 6 of 180 s. Stimuli were presented in 3 blocks with 4 stimuli in each block. In Study#2, two or three segments of degraded speech (7-Channels, 3-Channels; 4.8-21.6 s) were embedded in 15 clear audio streams. The lengths of the 15 stimuli varied between 60 s and 3 min with two stimuli of 60, 90, 120, and 9 of 180 s. Stimuli were presented in 5 blocks with 3 stimuli in each block. In both studies, each stimulus ended with a two-syllable noun within the last four words. In order to keep participants’ attention on the stimulation, we asked participants after each stimulus to choose from two presented two-syllable nouns, the one that had occurred within the last four words of a sentence. The sequence of all the audio stimuli was randomized across participants, not following the original storyline of the audiobook. The syllable rate of the stimuli varied between 3.1 and 4.3 Hz with a median of 4 Hz (estimated using Praat (*18*)).

### Vocoding

Noise-vocoding of all audio stimuli was done using the vocoder toolbox for MATLAB (*43*), and we created conditions with 7 and 3 channels (Fig. 1A). Vocoding for both studies was performed as described in (*11*). For the vocoding, the waveform of each audio stimulus was passed through two Butterworth analysis filters (for 7 and 3 channels) with a range of 200–7,000 Hz representing equal distances along the basilar membrane. Amplitude envelope extraction was done with half-wave rectification and low-pass filtering at 250 Hz. The envelopes were then normalized in each channel and multiplied with the carrier. Then, they were filtered in the band and the RMS of the resulting signal was adjusted to that of the original signal filtered in that same band. Auditory stimuli were presented binaurally using MEG-compatible pneumatic in-ear headphones (SOUNDPixx, VPixx technologies).

### Behavioral Assessment

Due to the low number of behavioral responses from the MEG part, we added an additional behavioral assessment. For Study#1 and Study#2, 24 audio files were created from recordings of another female native German speaker reading Antoiné St. Exupery’s “The little prince” (1943). Each stimulus contained one sentence (length between 2-15 s) and was either presented unvocoded with 7-channel vocoding or 3-channel vocoding. For Study #1, the stimuli in the vocoding condition were vocoded from start to the end; for Study #2, the stimuli were vocoded only in the last 0.6-5 s. Comparable to both MEG experiments, the stimuli also ended with a two-syllable noun within the last four words, and participants were asked to choose the last noun they heard between two nouns on the screen. The sequence of all audio stimuli was random across the participants, not following the storyline. In each study, the hit rates across the three vocoding conditions were compared using one-way repeated measures ANOVAs. Post-Hoc analysis was performed using FDR (*44*) corrected Wilcoxon signed-rank tests (as the assumptions for paired samples t-tests were violated).

### Data Acquisition

Data acquisition and parts of the data analysis for Study#1 and #2 closely resemble, with minor exceptions, the one described in two previous studies (*11, 45*). Magnetic brain activity was recorded using a 306-channel whole head MEG system (TRIUX, Elekta Oy, Finland) with a sampling rate of 1 kHz for the main experiments (Study#1 and Study#2) and with a sampling rate of 10 kHz for the brainstem localizer in Study#2 (see *Backward Modeling* for further information). The system consists of 204 planar gradiometers and 102 magnetometers. Before entering the magnetically shielded room (AK3B, Vakuumschmelze, Hanau, Germany), the head shape of each participant was acquired with >300 digitized points on the scalp, including fiducials (nasion, left and right pre-auricular points) with a Polhemus FASTRAK system (Polhemus, Vermont, USA). The auditory brainstem response was measured with a single electrode located on FpZ based on the electrode placement of the international 10–20-System (*46*). A ground electrode was placed on the forehead at midline and a reference on the clavicle bone of the participants.

## Data Analysis

### Preprocessing

All data analysis steps for Study #1 and #2 were performed similarly and are therefore reported together. The acquired data was Maxwell-filtered using a Signal Space Separation (SSS) algorithm (*47*) implemented in the Maxfilter program (version 2.2.15) provided by the MEG manufacturer to remove external magnetic interference from the MEG signal and realign data to a common standard head position (-trans default Maxfilter parameter). The Maxwell-filtered and continuous data was then further analysed using the FieldTrip toolbox (*48*) and custom built Matlab routines. First, the data was high-pass filtered at 0.1 Hz using a finite impulse response (FIR) filter (Kaiser window). For extracting physiological artefacts from the data, 50 independent components were calculated from the filtered data. Via visual inspection, the components showing eye-movements & heartbeats were removed from the data. On average across studies, 3 components were removed per subject (*SD* = 1). Then, trials related to each of the three conditions (Original, 7-Channels and 3-Channels) were defined. The acoustic speech envelope was extracted and aligned with the measured MEG data (*11*). Afterwards data was cut into segments of 4 seconds to increase signal-to-noise ratio.

### Source Analysis

Anatomical template images were warped to the individual head shape and brought into a common space by co-registering them based on the three anatomical landmarks (nasion, left and right preauricular points) with a standard brain from the Montreal Neurological Institute (MNI, Montreal, Canada) (*49*). Afterwards a single-shell head model (*50*) was computed for each participant. As a source model, a grid with 1 cm resolution and 2982 voxels based on an MNI template brain was morphed into the brain volume of each participant. This allows group-level averaging and statistical analysis as all the grid points in the warped grid belong to the same brain region across subjects. Common linearly constrained minimum variance (LCMV) beamformer spatial filters (*51*) were then computed on the preprocessed MEG data and applied to project the single-trial time series into source space. The number of epochs across conditions was equalized (by the lowest number of epochs across conditions within each study). We applied a frequency analysis to the 4-s segments of all three conditions (original, 7-Chan and 3-Chan) calculating multi-taper frequency transformation (dpss taper: 0–25 Hz in 0.25 Hz steps, 4 Hz smoothing, no baseline correction). For the coherence calculation between each virtual sensor and the acoustic speech envelope, 0.25-Hz frequency steps were chosen. Then, the coherence between activity at each virtual sensor and the acoustic speech envelope during acoustic stimulation in the frequency spectrum was calculated and averaged across trials. We refer to the coherence between acoustic speech envelope and brain activity as neural speech tracking. Most studies on neural speech tracking report findings of frequencies below 7 Hz; we, therefore, analysed frequencies between 2 and 7 Hz. We applied repeated-measures ANOVAs for each frequency within the range (*ft_statfun_depsamplesFunivariate* in FieldTrip) to test modulations of neural measures across the different intelligibility levels. To control for multiple comparisons, a nonparametric cluster-based permutation test test was undertaken (*52*). The test statistic was repeated 10,000 times on data shuffled across conditions and the largest statistical value of a cluster coherent in source space was kept in memory. The observed clusters were compared against the distribution obtained from the randomization procedure and were considered significant when their probability was below 5%. Effects were identified in source space. All voxels within the cluster and the corresponding individual coherence and power values were extracted and averaged. Post hoc paired samples *t* tests between conditions were corrected for multiple comparisons by using the FDR method (*44*) implemented in Pingouin (*53*). Slopes of the change in coherence along with changes in intelligibility were compared across studies using a Mann-Whitney *U* test. For visualization, source localizations were averaged across the 2–7 Hz frequency bands and mapped onto inflated surfaces as implemented in FieldTrip.

### Peak Analysis

For further analysis of the coherence spectra in source space, we extracted the most prominent peaks in the low frequency range (2-7Hz) across all virtual channels in which we observed a significant difference across vocoding levels (579 channels for Study#1 and 417 for Study#2; see Fig. 2 B, E). This was operationalized by using FOOOF (*12*) to flatten the coherence spectrum at each virtual channel and compute Gaussian model fits to extract peaks. For each subject, the average peak height, bandwidth and center frequency of the extracted peaks (see Fig. 3) were computed. Peaks were only considered if they exceeded a threshold relative to the aperiodic slope of 1.5 standard deviations (peak_threshold=1.5). Bad model fits were dropped (one bad model fit in Study#2). If the residual model fits differed from the rest based on the *R*^*2*^ (between the input spectrum and the full model fit), or error of the full model fit by more than 2.5 *SDs*, they were dropped. The most prominent peak in the range between 2 and 7 Hz was extracted per virtual channel. Peak and aperiodic parameters were then averaged across all virtual channels and further analysed using repeated-measures ANOVAs and dependent-samples *t*-tests (as implemented in Pingouin (*53*)) for post-hoc analysis (corrected for multiple comparisons using the FDR method (*44*)).

### Analysis of modulation- and syllable rate

We estimated the modulation and syllabic rate of all twelve audio files for each condition (Original, 7-Channels, 3-Channels). Audio files were transformed to 6-s duration segments (as in (*6*)) resulting in 386 audio segments per condition (Original, 7-Channels, 3-Channels). The modulation rates for the three different levels of intelligibility were then extracted using custom matlab scripts taken from (*6*). The center frequency of each spectrum was further extracted by taking the global maximum value of each modulation spectrum. The realized syllable rate of the speaker was computed using Praat (*18*). The center frequencies of the three conditions and syllable rate were then compared using repeated-measures ANOVAs and dependent-samples *t*-tests (as implemented in Pingouin (*53*)) for post-hoc analysis (corrected for multiple comparisons using the FDR method (*44*)). Afterwards an ensemble (50 classifiers) of *k*-nearest neighbor classifiers were trained in a nested 5-fold cross-validation (*54*) to decode whether a given frequency can be associated with either the modulation or the syllabic rate. We decided to use the *k*-nearest neighbor classifiers as data had only a low number of features (i.e. one center frequency per audio segment); a classification problem usually solved well by a *k*-nearest neighbor approach (*55*). The repeated nested cross-validation procedure was chosen to avoid overfitting of hyperparameters. Each external cross-validation loop was embedded in a repeated stratified *k*-folding procedure (*RepeatedStratifiedKFold;* 25 repetitions) the best number of neighbors was determined by searching the hyper-parameter space for the best cross-validation (CV) score of a kNN model using the implemented *GridSearchCV* function and by computing the area under the receiver operating characteristic curve (*roc-auc*) as loss-function. Confusion matrices were then computed on a separated test set (10% of all data) that was not part of the initial inner cross-validation to avoid overfitting of hyperparameters. Confusion matrices of each inner loop were kept in memory and averaged across all repetitions (150 repetitions). The procedure was implemented using sci-kit learn(*56*) and custom written python scripts. The code used for the analysis can be found in the corresponding authors gitlab repository (see Data & Code Availability). The trained classifiers were subsequently applied to the center frequencies from speech-brain coherence (see Peak Analysis) to determine whether a frequency was rather related to the modulation- or the syllabic rate. The corresponding probabilities were then compared using a two-way repeated measures ANOVA with the factors tracking (Modulation/Syllable rate) and vocoding (Original, 7-Channels, 3-Channels).

### Modeling of subcortical auditory activity

In order to reconstruct auditory brainstem activity from the MEG data we applied a recently developed backward modelling approach (*17*) to the data obtained in Study#2. As planar gradiometers are less sensitive to sources below the cortical surface than magnetometers (*57*) only magnetometer data was included in this analysis. The backward models were trained independently for each subject using data obtained from a localizer run dedicated to elicit auditory brainstem activity (see (*17*) for a detailed account). In brief, we used the signal captured by the MEG sensors (during the first 10ms) as regressors for a concurrent EEG recording of an auditory brainstem response (similar to the estimation of regression based ERPs (*58*)). The corresponding weights (a time-generalized representation of auditory brainstem activity) were then applied to the upsampled (10 000 Hz) single-trial time series data from Study#2. Afterwards, the data was downsampled (100Hz) and a frequency analysis was applied to the 4-s segments of all three conditions (original, 7-Chan and 3-Chan) calculating multi-taper frequency transformation (dpss taper: 0–25 Hz in 0.25 Hz steps, 4 Hz smoothing, no baseline correction) for the analyses of the coherence calculation between each virtual sensor and the acoustic speech envelope. Afterwards analysis steps that were performed for the previous analysis were repeated for the modeled activity (see statistics reported in source analysis and steps undertaken for peak and decoding analysis).

## Acknowledgements

This research was supported by the Sivantos GmbH and an FWF Einzelprojekt (P 31230).

## Data/Code Availability

The data and code necessary for generating the figures and computing statistics will be shared in the corresponding authors gitlab repository (https://gitlab.com/schmidtfa). Access to raw data will be made available upon reasonable request.

## Author contributions

Conceptualization: NW, AH, AK, FS, YC

Data curation: YC, FS, AH

Formal Analysis: FS, AH, YC

Funding acquisition: NW, AK, SR, RH, MS

Investigation: YC, FS, AH

Methodology: FS, NW

Project administration: NW

Software: FS, AH

Supervision: NW, RH, MS, AH, AK Visualization: FS

Writing—original draft: FS

Writing—review & editing: AH, AK, NW, YC, RH

## REFERENCES

1. J. Gross, N. Hoogenboom, G. Thut, P. Schyns, S. Panzeri, P. Belin, S. Garrod, Speech Rhythms and Multiplexed Oscillatory Sensory Coding in the Human Brain. PLoS Biol. 11, e1001752 (2013).

2. J. E. Peelle, J. Gross, M. H. Davis, Phase-Locked Responses to Speech in Human Auditory Cortex are Enhanced During Comprehension. Cereb. Cortex. 23, 1378–1387 (2013).

3. K. B. Doelling, L. H. Arnal, O. Ghitza, D. Poeppel, Acoustic landmarks drive delta–theta oscillations to enable speech comprehension by facilitating perceptual parsing. NeuroImage. 85, 761–768 (2014).

4. A. Keitel, J. Gross, C. Kayser, Perceptually relevant speech tracking in auditory and motor cortex reflects distinct linguistic features. PLOS Biol. 16, e2004473 (2018).

5. D. Poeppel, M. F. Assaneo, Speech rhythms and their neural foundations. Nat. Rev. Neurosci. 21, 322–334 (2020).

6. N. Ding, A. D. Patel, L. Chen, H. Butler, C. Luo, D. Poeppel, Temporal modulations in speech and music. Neurosci. Biobehav. Rev. 81, 181–187 (2017).

7. C. Coupé, Y. Oh, D. Dediu, F. Pellegrino, Different languages, similar encoding efficiency: Comparable information rates across the human communicative niche. Sci. Adv. 5, eaaw2594 (2019).

8. E. Jacewicz, R. A. Fox, C. O’Neill, J. Salmons, Articulation rate across dialect, age, and gender. Lang. Var. Change. 21, 233–256 (2009).

9. N. Ding, M. Chatterjee, J. Z. Simon, Robust cortical entrainment to the speech envelope relies on the spectro-temporal fine structure. NeuroImage. 88, 41–46 (2014).

10. J. Song, P. Iverson, Listening effort during speech perception enhances auditory and lexical processing for non-native listeners and accents. Cognition. 179, 163–170 (2018).

11. A. Hauswald, A. Keitel, Y. Chen, S. Rösch, N. Weisz, Eur. J. Neurosci., in press, doi:10.1111/ejn.14912.

12. T. Donoghue, M. Haller, E. J. Peterson, P. Varma, P. Sebastian, R. Gao, T. Noto, A. H. Lara, J. D. Wallis, R. T. Knight, A. Shestyuk, B. Voytek, Parameterizing neural power spectra into periodic and aperiodic components. Nat. Neurosci. 23, 1655–1665 (2020).

13. R. F. Voss, J. Clarke, ‘1/fnoise’ in music and speech. Nature. 258, 317–318 (1975).

14. W. S. Pritchard, The Brain in Fractal Time: 1/F-Like Power Spectrum Scaling of the Human Electroencephalogram. Int. J. Neurosci. 66, 119–129 (1992).

15. H. Wen, Z. Liu, Separating Fractal and Oscillatory Components in the Power Spectrum of Neurophysiological Signal. Brain Topogr. 29, 13–26 (2016).

16. R. V. Shannon, F.-G. Zeng, V. Kamath, J. Wygonski, M. Ekelid, Speech Recognition with Primarily Temporal Cues. Science. 270, 303–304 (1995).

17. F. Schmidt, G. Demarchi, F. Geyer, N. Weisz, A backward encoding approach to recover subcortical auditory activity. NeuroImage. 218, 116961 (2020).

18. N. H. de Jong, T. Wempe, Praat script to detect syllable nuclei and measure speech rate automatically. Behav. Res. Methods. 41, 385–390 (2009).

19. A. E. Forte, O. Etard, T. Reichenbach, The human auditory brainstem response to running speech reveals a subcortical mechanism for selective attention, 12.

20. O. Etard, M. Kegler, C. Braiman, A. E. Forte, T. Reichenbach, Decoding of selective attention to continuous speech from the human auditory brainstem response. NeuroImage. 200, 1–11 (2019).

21. R. K. Maddox, A. K. C. Lee, eneuro, in press, doi:10.1523/ENEURO.0441-17.2018.

22. M. J. Polonenko, R. K. Maddox, Exposing distinct subcortical components of the auditory brainstem response evoked by continuous naturalistic speech. eLife. 10, e62329 (2021).

23. Q. Gehmacher, P. Reisinger, T. Hartmann, T. Keintzel, S. Rösch, K. Schwarz, N. Weisz, “Direct cochlear recordings show human hearing nerve activity is modulated by selective attention” (preprint, Neuroscience, 2021),, doi:10.1101/2021.03.01.433316.

24. M. H. Albrecht Köhler, G. Demarchi, N. Weisz, “Cochlear activity in silent cue-target intervals shows a theta-rhythmic pattern and is correlated to attentional alpha modulations” (preprint, Neuroscience, 2019),, doi:10.1101/653311.

25. K. von Kriegstein, R. D. Patterson, T. D. Griffiths, Task-Dependent Modulation of Medial Geniculate Body Is Behaviorally Relevant for Speech Recognition. Curr. Biol. 18, 1855–1859 (2008).

26. H. Luo, D. Poeppel, Phase Patterns of Neuronal Responses Reliably Discriminate Speech in Human Auditory Cortex. Neuron. 54, 1001–1010 (2007).

27. L. Decruy, J. Vanthornhout, T. Francart, Hearing impairment is associated with enhanced neural tracking of the speech envelope. Hear. Res. 393, 107961 (2020).

28. A. Presacco, J. Z. Simon, S. Anderson, Speech-in-noise representation in the aging midbrain and cortex: Effects of hearing loss. PLOS ONE. 14, e0213899 (2019).

29. B. Mirkovic, S. Debener, J. Schmidt, M. Jaeger, T. Neher, Effects of directional sound processing and listener’s motivation on EEG responses to continuous noisy speech: Do normal-hearing and aided hearing-impaired listeners differ? Hear. Res. 377, 260–270 (2019).

30. R. Reetzke, G. N. Gnanateja, B. Chandrasekaran, Neural tracking of the speech envelope is differentially modulated by attention and language experience. Brain Lang. 213, 104891 (2021).

31. N. Molinaro, M. Lizarazu, M. Lallier, M. Bourguignon, M. Carreiras, Out-of-synchrony speech entrainment in developmental dyslexia: Altered Cortical Speech Tracking in Dyslexia. Hum. Brain Mapp. 37, 2767–2783 (2016).

32. J. Ahveninen, I. P. Jaaskelainen, T. Raij, G. Bonmassar, S. Devore, M. Hamalainen, S. Levanen, F.-H. Lin, M. Sams, B. G. Shinn-Cunningham, T. Witzel, J. W. Belliveau, Task-modulated “what” and “where” pathways in human auditory cortex. Proc. Natl. Acad. Sci. 103, 14608–14613 (2006).

33. S. Atiani, M. Elhilali, S. V. David, J. B. Fritz, S. A. Shamma, Task Difficulty and Performance Induce Diverse Adaptive Patterns in Gain and Shape of Primary Auditory Cortical Receptive Fields. Neuron. 61, 467–480 (2009).

34. S. Atiani, S. V. David, D. Elgueda, M. Locastro, S. Radtke-Schuller, S. A. Shamma, J. B. Fritz, Emergent Selectivity for Task-Relevant Stimuli in Higher-Order Auditory Cortex. Neuron. 82, 486–499 (2014).

35. C. Demanuele, C. J. James, E. J. Sonuga-Barke, Distinguishing low frequency oscillations within the 1/f spectral behaviour of electromagnetic brain signals. Behav. Brain Funct. 3, 62 (2007).

36. B. Voytek, M. A. Kramer, J. Case, K. Q. Lepage, Z. R. Tempesta, R. T. Knight, A. Gazzaley, Age-Related Changes in 1/f Neural Electrophysiological Noise. J. Neurosci. 35, 13257–13265 (2015).

37. M. M. Robertson, S. Furlong, B. Voytek, T. Donoghue, C. A. Boettiger, M. A. Sheridan, EEG power spectral slope differs by ADHD status and stimulant medication exposure in early childhood. J. Neurophysiol. 122, 2427–2437 (2019).

38. J. L. Molina, B. Voytek, M. L. Thomas, Y. B. Joshi, S. G. Bhakta, J. A. Talledo, N. R. Swerdlow, G. A. Light, Memantine Effects on Electroencephalographic Measures of Putative Excitatory/Inhibitory Balance in Schizophrenia. Biol. Psychiatry Cogn. Neurosci. Neuroimaging. 5, 562–568 (2020).

39. B. J. He, J. M. Zempel, A. Z. Snyder, M. E. Raichle, The Temporal Structures and Functional Significance of Scale-free Brain Activity. Neuron. 66, 353–369 (2010).

40. E. Podvalny, N. Noy, M. Harel, S. Bickel, G. Chechik, C. E. Schroeder, A. D. Mehta, M. Tsodyks, R. Malach, A unifying principle underlying the extracellular field potential spectral responses in the human cortex. J. Neurophysiol. 114, 505–519 (2015).

41. N. Ding, L. Melloni, H. Zhang, X. Tian, D. Poeppel, Cortical tracking of hierarchical linguistic structures in connected speech. Nat. Neurosci. 19, 158–164 (2016).

42. B. Diaz, F. Hintz, S. J. Kiebel, K. von Kriegstein, Dysfunction of the auditory thalamus in developmental dyslexia. Proc. Natl. Acad. Sci. 109, 13841–13846 (2012).

43. E. Gaudrain, Vocoder: Basal (Zenodo, 2016; https://zenodo.org/record/48120).

44. Y. Benjamini, Y. Hochberg, Controlling the False Discovery Rate: A Practical and Powerful Approach to Multiple Testing. J. R. Stat. Soc. Ser. B Methodol. 57, 289–300 (1995).

45. A. Hauswald, C. Lithari, O. Collignon, E. Leonardelli, N. Weisz, A Visual Cortical Network for Deriving Phonological Information from Intelligible Lip Movements. Curr. Biol. 28, 1453-1459.e3 (2018).

46. G. H. Klem, H. O. Lüders, H. H. Jasper, C. Elger, The ten-twenty electrode system of the International Federation. The International Federation of Clinical Neurophysiology. Electroencephalogr. Clin. Neurophysiol. Suppl. 52, 3–6 (1999).

47. S. Taulu, J. Simola, Spatiotemporal signal space separation method for rejecting nearby interference in MEG measurements. Phys. Med. Biol. 51, 1759–1768 (2006).

48. R. Oostenveld, P. Fries, E. Maris, J.-M. Schoffelen, FieldTrip: Open Source Software for Advanced Analysis of MEG, EEG, and Invasive Electrophysiological Data. Comput. Intell. Neurosci. 2011, 1–9 (2011).

49. J. Mattout, R. N. Henson, K. J. Friston, Canonical Source Reconstruction for MEG. Comput. Intell. Neurosci. 2007, 1–10 (2007).

50. G. Nolte, The magnetic lead field theorem in the quasi-static approximation and its use for magnetoencephalography forward calculation in realistic volume conductors. Phys. Med. Biol. 48, 3637–3652 (2003).

51. B. D. Van Veen, W. Van Drongelen, M. Yuchtman, A. Suzuki, Localization of brain electrical activity via linearly constrained minimum variance spatial filtering. IEEE Trans. Biomed. Eng. 44, 867–880 (1997).

52. E. Maris, R. Oostenveld, Nonparametric statistical testing of EEG-and MEG-data. J. Neurosci. Methods. 164, 177–190 (2007).

53. R. Vallat, Pingouin: statistics in Python. J. Open Source Softw. 3, 1026 (2018).

54. M. Hosseini, M. Powell, J. Collins, C. Callahan-Flintoft, W. Jones, H. Bowman, B. Wyble, I tried a bunch of things: The dangers of unexpected overfitting in classification of brain data. Neurosci. Biobehav. Rev. 119, 456–467 (2020).

55. D. A. Eisa, A. I. Taloba, S. S. I. Ismail, A Comparative Study on using Principle Component Analysis with different Text Classifiers. Int. J. Comput. Appl. 180, 1–6 (2018).

56. F. Pedregosa, G. Varoquaux, A. Gramfort, V. Michel, B. Thirion, O. Grisel, M. Blondel, P. Prettenhofer, R. Weiss, V. Dubourg, J. Vanderplas, A. Passos, D. Cournapeau, M. Brucher, M. Perrot, É. Duchesnay, Scikit-learn: Machine Learning in Python. J. Mach. Learn. Res. 12, 2825–2830 (2011).

57. J. Vrba, S. E. Robinson, SQUID sensor array configurations for magnetoencephalography applications. Supercond. Sci. Technol. 15, R51–R89 (2002).

58. N. J. Smith, M. Kutas, Regression-based estimation of ERP waveforms: I. The rERP framework: rERPs I. Psychophysiology. 52, 157–168 (2015).

59. J. Z. Bakdash, L.R. Marusich, Repeated Measures Correlation. Front. Psychol. 8 (2017), doi:10.3389/fpsyg.2017.00456.

