## Supplementary Material for "Neural speech tracking shifts from the syllabic to the modulation rate of speech as intelligibility decreases"

### SUPPLEMENTARY MATERIALS

**Text S1. Decreases in intelligibility can be associated with a lower offset and flatter slope of low frequency speech-brain coherence**

In this study, we mainly focused on investigating the periodic components underlying speech-brain coherence. However, when separating the periodic from the aperiodic components we also noticed that exponent and offset of speech-brain coherence were modulated by intelligibility (see Fig. S1). We further investigated this using a repeated-measure ANOVA. This analysis showed that in both studies the exponent (Study#1, $F(2, 54) = 45.898$, $p_{ggeisser} = 2.329e^{-10}$, $\eta_p^2 = 0.630$; Study#2, $F(2, 50) = 18.409$, $p = 1e^{-06}$, $\eta_p^2 = 0.424$) and the offset (Study#1, $F(2, 54) = 51.793$, $p = 2.765e^{-13}$, $\eta_p^2 = 0.657$; Study#2, $F(2, 50) = 35.804$, $p = 2.239e^{-10}$, $\eta_p^2 = 0.589$) differed significantly across the three vocoding conditions. Comparing the different vocoding levels with each other showed a higher coherence offset for unvocoded stimuli than for stimuli vocoded with 7-Channels (Study#1, $t(27) = 3.505$, $p_{fdr} = 0.0016$, $d = 0.744$; Study#2, $t(25) = 2.214$, $p_{fdr} = 0.0361$, $d = 0.466$) and with 3-Channels (Study#1, $t(27) = 8.289$, $p_{fdr} = 1.598e^{-08}$, $d = 2.156$; Study#2, $t(25) = 8.581$, $p_{fdr} = 1.919e^{-08}$, $d = 2.031$). Additionally stimuli vocoded with 7-Channels had a higher offset than stimuli vocoded with 3-Channels (Study#1, $t(27) = 7.559$, $p_{fdr} = 5.912e^{-08}$, $d = 1.486$; Study#2, $t(25) = 5.676$, $p_{fdr} = 6.558e^{-06}$, $d = 1.277$).

Furthermore, the exponent of the slope of the coherence spectra flattened with intelligibility. Unvocoded stimuli elicited a steeper slope in the coherence spectra than stimuli vocoded with 7-Channels (in Study#1; $t(27) = 3.841$, $p_{fdr} = 0.0007$, $d = 0.936$, but not in Study#2 ($t(25) = 1.268$, $p_{fdr} = 0.216$, $d = 0.259$) and stimuli vocoded with 3-Channels (Study#1, $t(27) = 7.961$, $p_{fdr} = 4.431e^{-08}$, $d = 2.304$; Study#2, $t(25) = 6.171$, $p_{fdr} = 6e^{-06}$, $d = 1.488$). Additionally stimuli vocoded with 7-Channels elicited a steeper slope than stimuli vocoded with 3-Channels (Study#1, $t(27) = 7.528$, $p_{fdr} = 6.377e^{-08}$, $d = 1.246$; Study#2, $t(25) = 4.047$, $p_{fdr} = 0.0007$, $d = 0.993$).

In sum, we show that as intelligibility decreases the offset of speech-brain coherence decreases and the slope of the coherence spectra becomes flatter. These differences in the aperiodic parameters may also partly account for band-limited differences in low frequency speech-brain coherence that are commonly related and also interpreted as differences in neural speech tracking. Yet, the separation of broadband coherence in periodic and aperiodic components illustrates that analysing coherence in a band-limited range, even though more or less explicitly assumed, may not reflect neural tracking of sound or linguistic information in the relevant frequency range. But to what extent are these parameters related to the band-limited differences in low frequency speech tracking that we measured (see Fig. 2)?

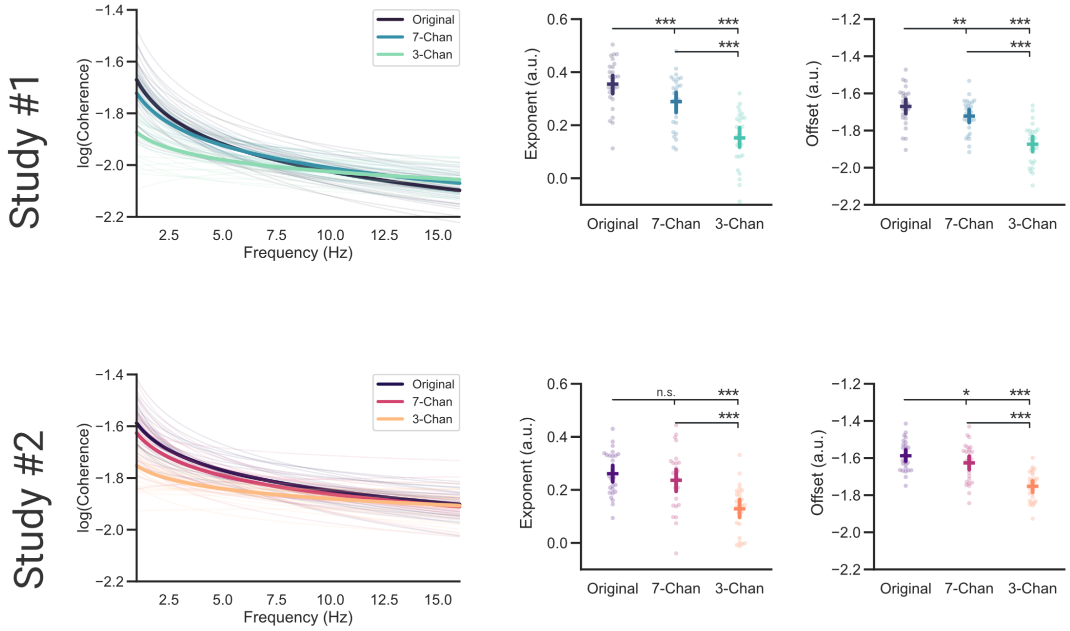


***Fig. S1 Decreases in intelligibility can be associated with a lower offset and flatter slope of low frequency speech-brain coherence.*** *The averaged exponents and offsets extracted from the coherence spectra for each subject were compared across three conditions (Original, 7-Chan & 3-Chan). Bars represent 95% confidence intervals,* $p_{fdr} < 0.05$*, $p_{fdr} < 0.01$**, $p_{fdr} < 0.001$***

### Text S2. Aperiodic components explain most of the variance of low frequency speech-brain coherence

In order to better understand the extent to which aperiodic components explain the effect we showed for low frequency speech-brain coherence (see Fig. 2) we calculated a repeated measures correlation (*59*) between the averaged coherence in the low frequency range (2-7Hz) and the periodic and aperiodic components in the same frequency range across all vocoding levels (Original, 7-Channels, 3-Channels). The results of this analysis showed that low-frequency speech-brain coherence showed overall higher correlation coefficients with the underlying aperiodic components offset (Study#1, $r$ (55) = 0.91, $p$ = 8.476e$^{-23}$, $r^2$ = 0.83; Study#2, $r$ (51) = 0.6, $p$ = 2e$^{-06}$, $r^2$ = 0.36) and exponent (Study#1, $r$ (55) = 0.82, $p$ = 8.726e$^{-15}$, $r^2$ = 0.67; Study#2, $r$ (51) = 0.57, $p$ = 1e$^{-05}$, $r^2$ = 0.32) compared to the periodic components center frequency (Study#1, $r$ (55) = -0.56, $p$ = 7e$^{-06}$, $r^2$ = 0.31; Study#2, $r$ (51) =

-0.22, *p* = 0.121, $r^2$ = 0.05), bandwidth (Study#1, *r* (55) = 0.7, *p* = 1.948e$^{-09}$, $r^2$ = 0.48; Study#2, *r* (51) = 0.41, *p* = 0.002, $r^2$ = 0.17) and the relative magnitude of the coherence peak (Study#1, *r* (55) = 0.33, *p* = 0.011, $r^2$ = 0.11; Study#2, *r* (51) = 0.00, *p* = 0.998, $r^2$ = 0.00).

In sum, these findings illustrate the importance of parametrizing speech-brain coherence spectra to remove the influence of aperiodic components from the periodic components to better understand the parameters that actually reflect neural speech tracking.

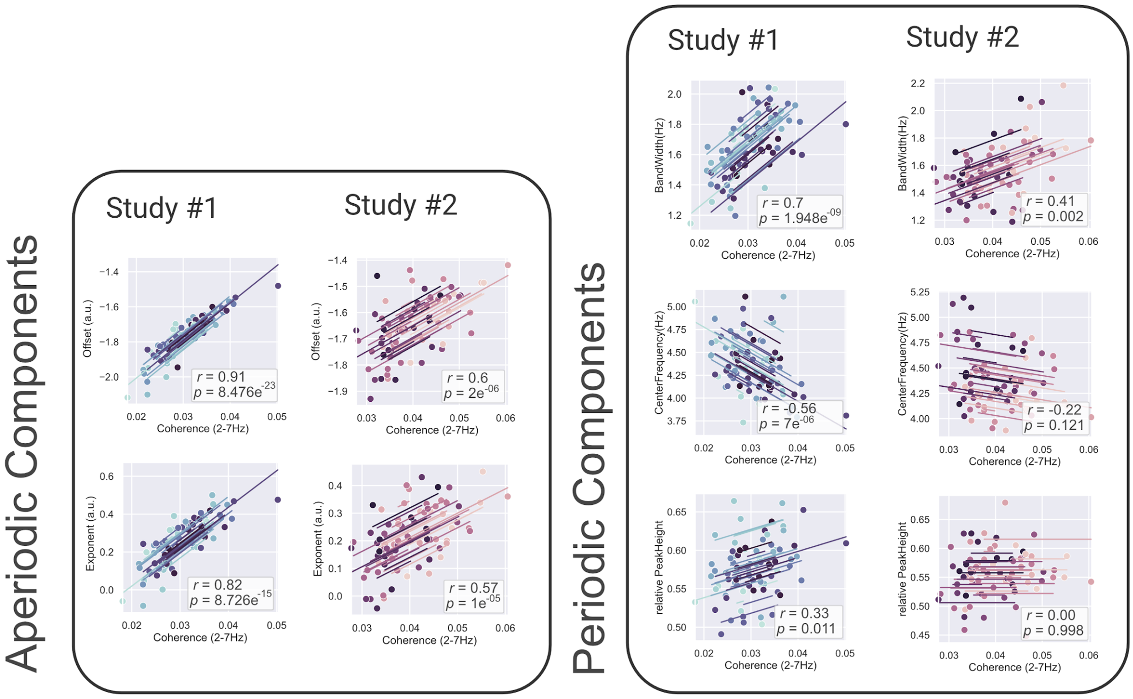


***Fig. S2 Aperiodic components explain most of the variance of low frequency speech- brain coherence***. *Using a repeated measures correlation analysis we show that the aperiodic components (offset, exponent) of speech-brain coherence were related stronger to the band-limited (2-7Hz) averaged speech-brain coherence than the periodic components (center frequency, relative magnitude, bandwidth). This highlights the importance of parametrizing speech-brain coherence to better understand neural speech tracking.*

**Table S1. Differences across the extracted center frequencies and the syllable rate of the audio signal.** Post-Hoc analysis was performed using $fdr_{bh}$ corrected paired samples wilcoxon signed-rank tests implemented in pingouin ***(53)***

| A | B | W-val | $p_{fdr}$ | Cohen's *d* |
|---|---|---|---|---|
| CF 3-Chan | CF 7-Chan | 2730.5 | $8.847e^{-12}$ | -0.345 |
| CF 3-Chan | CF Original | 3688.5 | $5.478e^{-26}$ | -0.842 |
| CF 3-Chan | SyllableRate | 2642.5 | $5.271e^{-52}$ | 1.640 |
| CF 7-Chan | CF Original | 9390.5 | $1.716e^{-10}$ | -0.433 |
| CF 7-Chan | SyllableRate | 1591.0 | $1.022e^{-55}$ | 1.892 |
| CF Original | SyllableRate | 177.5 | $1.705e^{-60}$ | 2.747 |
